# The relationship between heart rate variability and cerebral autoregulation at elevated intracranial pressure

**DOI:** 10.64898/2026.11.25.769098

**Authors:** Zeynep Ozkaya, Jingyi Wu, Jana Kainerstorfer

## Abstract

**Objective:** Altered intracranial pressure (ICP) is a key biomarker for traumatic brain injury (TBI). However, obtaining ICP measurements is a highly invasive process. Heart rate variability (HRV) has demonstrated potential as a useful biomarker for ICP changes, but few studies have investigated the relationship between ICP changes and HRV. This work examines how HRV and ICP interact in cases of TBI and whether HRV metrics can be informative of the state of autoregulatory control at elevated ICP.

**Approach:** Electrocardiogram signals were recorded in seven non-human primates under anesthesia. ICP changes were induced by means of fluid increase via a cannula in the ventricles. Time- and frequency-domain HRV metrics were calculated from ECG and correlated with physiological indicators including CPP, ICP, and pressure reactivity index (PRx).

**Main results:** Results demonstrate that there is no direct linear relationship between HRV and ICP. PRx and HRV, however, revealed a significant group level relationship where the standard deviation of normal-to-normal intervals (SDNN) increased after a clinically significant PRx threshold, implying that HRV may be more closely associated with TBIs in terms of autoregulatory changes rather than ICP changes. . This study provides insights into changes in brain physiology in response to ICP increases by demonstrating the relationship between CAR and HRV. Further, this work draws attention to conflicting reports in the literature when interpreting the interaction between HRV and other physiological variables.

## 1. Introduction

Intracranial pressure (ICP) is a key biomarker for traumatic brain injury (TBI), where severe impact to the skull can cause physiological changes such as local tissue damage and obstruction to blood and cerebrospinal fluid (CSF) flow (Smith 2008, Hawryluk *et al* 2022). These TBI induced brain tissue and fluid volume changes can cause an increase in the pressure within skull and lead to pathological outcomes such as cerebral ischemia or intracranial hypertension (Jha *et al* 2019, Hlatky *et al* 2003). Thus, ICP monitoring is crucial for assessing the state of brain health following TBI. However, standard ICP measurement methods are highly invasive, involving insertion of a catheter or micro transducer devices (Raboel *et al* 2012). Non-invasive techniques for ICP monitoring, such as diffuse correlational spectroscopy or transcranial doppler, have shown promising results in *in vivo* testing, but have yet to be translated to clinical settings as replacements of gold standard invasive techniques (Ruesch *et al* 2020, Müller *et al* 2023).

Fluctuations in ICP, however, can induce other physiological effects that can be used to assess the state of cerebral health based on clinically available signals. Notably, because raised ICP can obstruct cerebral blood flow (CBF), cerebral autoregulation (CAR), which is the process by which the cerebral vasculature modulates blood flow, is also impacted. Beyond cerebral autoregulation, raised ICP from TBI may disrupt autonomic regulation of the cardiovascular system. Thus, physiological signals beyond the cerebral vasculature can provide additional information about cerebral health (Paulson *et al* 1990, Hlatky *et al* 2003, Depreitere *et al* 2014). Heart rate variability (HRV), in particular, has shown promise for monitoring TBIs and can be used to predict long-term outcomes for patients (Pinto *et al* 2024, Sykora *et al* 2016). HRV is defined as the variability in the time intervals between adjacent heart beats and can be calculated using both time- and frequency-domain methods. The “gold-standard” time-domain measure is the standard deviation of the inter-beat interval (IBI) of normal sinus beats (SDNN), which directly measures the variability in the time intervals between successive heartbeat peaks.

Variations in these IBIs have been attributed to parasympathetic mediation of the heart by directly linking parasympathetic cardiac control to the degree of physiologically healthy heart rate variations (Katona and Jih 1975). Frequency-domain metrics have also shown correlations with nervous system activity, with high frequency (HF) HRV reflecting parasympathetic nervous system (PNS) activity. No causal link, however, has been made between the sympathetic nervous system (SNS) and HRV metrics, although sympathetic mediation of the heartbeats can lead to more rapid and regular timing of beats (Shaffer and Ginsberg 2017, Shaffer *et al* 2014, Goldstein *et al* 2011). Low frequency (LF) HRV has been related to both the PNS and SNS, but most closely tracks the baroreflex activity where slow oscillations are indicative of the baroreflex’s adjustment of the heart rate in response to changes in blood pressure. Higher HRV could also reflect autonomic flexibility while lower HRV reflects a less adaptive nervous system, corresponding to worse outcomes (Friedman 2007). However, this linear relation does not always hold in clinical settings and must be treated in a context dependent and dynamical manner. Further, many studies report contradicting results, with no clear consensus on whether elevated or reduced HRV corresponds to dangerous physiological conditions (Pinto *et al* 2024, Biswas *et al* 2000, Laborde *et al* 2017, Ghazzal *et al* 2026). Thus, it remains debated how exactly HRV is linked to changes in cerebral vasculature at times of elevated ICP.

This gap, along with the close interplay among ICP, cerebral perfusion, hemodynamics, and HRV in TBI, highlights the need to better understand the physiological mechanisms driving the heart’s response to changes in brain physiology. To address this, this work examines whether non-invasively measured HRV metrics can be informative of the state of CAR and autonomic control in cases of elevated ICP. Time- and frequency-domain HRV metrics were first correlated with systematically elevated ICP levels, revealing no direct linear relationship. However, further analysis of the relationship between HRV and CAR metrics suggests that HRV can be informative of the state of autoregulation at elevated ICP, consistent with observed interactions between cardiac autonomic activity and cerebrovascular control (Mowery *et al* 2008, Fedriga *et al* 2021).

## 2. Methods

Changes in fluid volume were induced in non-human primates (NHP) to investigate how elevated ICP correlates with CAR and HRV during simultaneous recording of pressure and heart signals. SDNN, root mean square of successive RR intervals (RMSSD), normalized LF power, normalized HF power, and the ratio between LF and HF components of IBI (LF/HF Ratio) were selected as time- and frequency-domain HRV metrics. These metrics were calculated from electrocardiogram (ECG) signals. SDNN is the most direct measure of changes in HRV, which is influenced by both SNS and PNS activation and RMSSD is a time invariant measure of beat-to-beat variance. The LF and HF metrics alone provide information about the SNS and vagal activity independently, while the LF/HF ratio can provide insight into the relative contributions of the SNS and PNS. Normalized LF and HF values were calculated to highlight relative changes over absolute values.

### 2.1 Data Collection

All experimental procedures complied with guidelines in the National Institute of Health’s Guide for the Care and Use of Laboratory Animals and were approved by the University of Pittsburgh and Carnegie Mellon University’s Institutional Animal Care and Use Committees. Data on 7 anesthetized NHPs (NHPs 1 to 3 Macaca mulatta aged 7.9 ± 1.5 years, weighing 9.4 ± 0.7 kg and NHPs 4 to 7 Macaca fascicularis aged 4.2 ± 0.9 years, weighing 5.1 ± 2.1 kg) was collected while changing ICP levels. Each NHP was sedated with either 20-mg/kg of ketamine or a combination of 1 mg/kg Diazepam and 0.04 mg/kg of Atropine before being transported to the experimentation room. The NHPs were anesthetized with 10 to 25.6 μg⁄kg⁄h fentanyl, and a small amount of isoflurane gas (< 1%) during the experiments and 0.1-mg⁄kg⁄h of vecuronium bromide paralytic was administered intravenously.

Heart rate, respiration rate, arterial blood oxygenation (SaO2), end tidal carbon dioxide partial pressure (ETCO2), and other vital signs were monitored using the SurgiVet veterinary monitor (Smiths Medical, Minneapolis, MN, USA). The NHPs were ventilated at 0.18-0.4 Hz. Two small craniotomies were made for monitoring ICP and inducing pressure changes. A catheter (Lumbar catheter, Medtronic, Minneapolis MN) was placed on the lateral ventricle through one of the craniotomies. An ICP sensor (Precision Pressure Catheter, Raumedic Helmbrechts, Germany) was placed on the brain parenchyma through the second craniotomy. The catheter placed on the lateral ventricle was connected to a saline reservoir, which was used to vary the ICP. Changes in the height of the saline reservoir led to fluid flow into the ventricle, which induced an ICP change. Arterial blood pressure (ABP) was measured through an arterial line and was measured with a sampling frequency of 100 Hz using a MPR1 Datalogger (Raumedic Helmbrechts, Germany). The ICP sensor as well as the parenchymal pressure sensor were connected to the MPR1 Datalogger and sampled at a rate of 100 Hz. In NHPs 1-3, a 3-lead ECG was placed on the chest with data recorded using an in-house amplifier circuit that registered a cardiac pulse when the ECG signal exceeded an empirically determined threshold for the R peak of the QRS complex. A synchronization pulse was then sent to an external auxiliary port. This cardiac pulse measurement was sampled at 100 Hz. In NHPs 4-7, ECG signal was measured using a separate device sampled at 1000 Hz. A schematic of the experimental set up, as well as representative time-traces are plotted in Figure 1.

**Figure 1.**
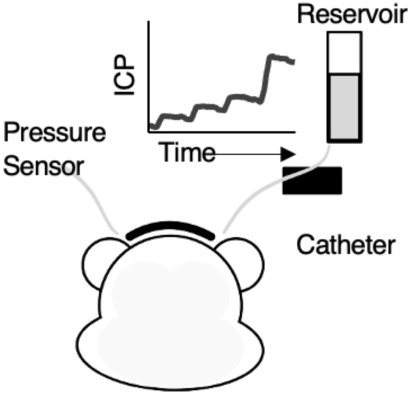
Schematic of experimental set-up with pressure sensor, saline reservoir, and catheter placement and representative plot of ICP increase over time.

Each experiment lasted 16.2 hours on average (SD 5.1 h, range 10.25 to 24.5 h). Each study contained 6-10 periods in which ICP was changed from the baseline using the saline reservoir. In NHPs 1-3, ICP increases were gradually induced between 9 mmHg and 30 mmHg. In NHPs 4-7, ICP increases were gradually induced between 5 mmHg and 60 mmHg. For all subjects baseline recording of ICP was taken before ICP was varied and each trial lasted approximately 90 minutes (Relander *et al* 2022, Ruesch *et al* 2020).

### 2.2 Inter-beat Interval

Both time- and frequency-domain HRV metrics are derived from the IBI (Shaffer and Ginsberg 2017). For NHPs 1-3, outliers were removed from the raw ECG trace on a point-by-point basis by first computing a median around a small window. Then, an outlier was flagged if the point surpassed an empirically determined threshold. Following outlier removal, the data was further visually inspected to confirm adequate artifact removal. Because the ECG signal for NHPs 1-3 contain only the cardiac pulses, the IBI was calculated by taking the difference of this signal to determine the time difference between the pulses. The discrete IBI trace was then smoothed in a 25 second window for additional cleaning.

The raw ECG signal for NHPs 4-7 was first bandpass filtered between 0.2 and 40 Hz using a fourth-order zero-phase Butterworth filter before outlier removal was performed as in NHPs 1-3. An additional smoothing filter was then applied with a window of 50 seconds. Then, cardiac peaks were detected using the *find_peaks* function form the SciPy signal-processing library to locate the point at which each QRS peak occurred (Virtanen *et al* 2020). Following peak detection, the IBI was computed in the same manner as NHPs 1-3. Outliers were removed from the extracted IBI for all NHPs on a point-by-point basis relative to the small window mean and then the signals were interpolated to 100 Hz using the *CubicSpline* function from the SciPy library to align with the time values for the pressure variables of ABP, ICP, and cerebral perfusion pressure (CPP).

### 2.3 Heart Rate Variability

The SDNN was calculated through a moving standard deviation of the IBI in 60-second bins, each with 45 seconds of overlap over the entire duration of the experiment. A 60-second window was used rather than the traditional 5-minute window because the recording only spanned one session rather than multiple days, which is more typical in clinical contexts (Shaffer and Ginsberg 2017). The RMSSD was calculated by taking the root mean square in this same window as a measure of variance in the heartbeats. The pressure variables, after undergoing the same outlier removal and smoothing process as the ECG signals, were aligned in time to the ECG and averaged within the same moving window to maintain alignment.

For the frequency domain HRV calculations, the IBI signals were first detrended to account for slow drift in the signal. Then, the Fast-Fourier Transform (FFT) of the IBI was calculated within the same 60-second window as time-domain HRV and the pressure averages. LF power was taken from the 0.04Hz to 0.15Hz band of the IBI and HF power was taken from 0.15Hz to 0.4Hz. LF and HF values were normalized by dividing by the combined LF + HF. The LF/HF ratio was calculated from these values (Shaffer and Ginsberg 2017).

### 2.4 Pressure Reactivity Index

In order to estimate CAR over fluctuating ICP, pressure reactivity index (PRx), which is the correlation between ABP and ICP, was calculated within the same 60-second bins. While standard clinical PRx is calculated in 5-minute windows, the shorter time scales and higher sampling rate of the data allowed for the use of shorter analysis windows as in the HRV calculations. PRx is indicative of the state of CAR, reflecting how closely fluctuations in blood pressure are related to ICP, with a weaker relationship expected when the autoregulatory response is intact. Although there is no true threshold, most clinical reports show a correlation >.3 indicating impaired autoregulation, while a negative or low coefficient implies intact autoregulation (Czosnyka *et al* 2017).

## 3. Results

Results indicate that HRV is informative of the state of autoregulation but has no significant relationship with ICP at the group level. While HRV metrics alone were insufficient for monitoring elevations in ICP, changes in HRV were informative of increases in ICP during points at which CAR was disrupted. HRV metrics tracked autoregulatory changes at elevated ICP levels with significance at subject level. Group level analysis demonstrates similar overall trends, though without statistical significance, suggesting the lack of a direct linear relationship between HRV and ICP. However, comparison of HRV metrics above and below the clinically relevant PRx threshold showed a significant difference in SDNN, suggesting that HRV may be informative of changes in autoregulatory state.

### 3.1 Time Domain HRV Metrics Inconsistently Track ICP Fluctuations

Figure 2 plots the pressure values, along with the SDNN and IBI of each subject over the entire time course of each session. Here, it is evident that at points of fluctuating ICP, for instance, between 500-600 minutes for subject 3 or 50-150 minutes for subject 4, the SDNN tracks the changes in ICP, ABP, CPP. These points in time at which HRV follows ICP fluctuations, however, are not consistent across subjects in terms of frequency, ICP level, or duration. In addition, in cases where ICP is more stable, the SDNN time trace reflects this, as seen with subject 5.

**Figure 2.**
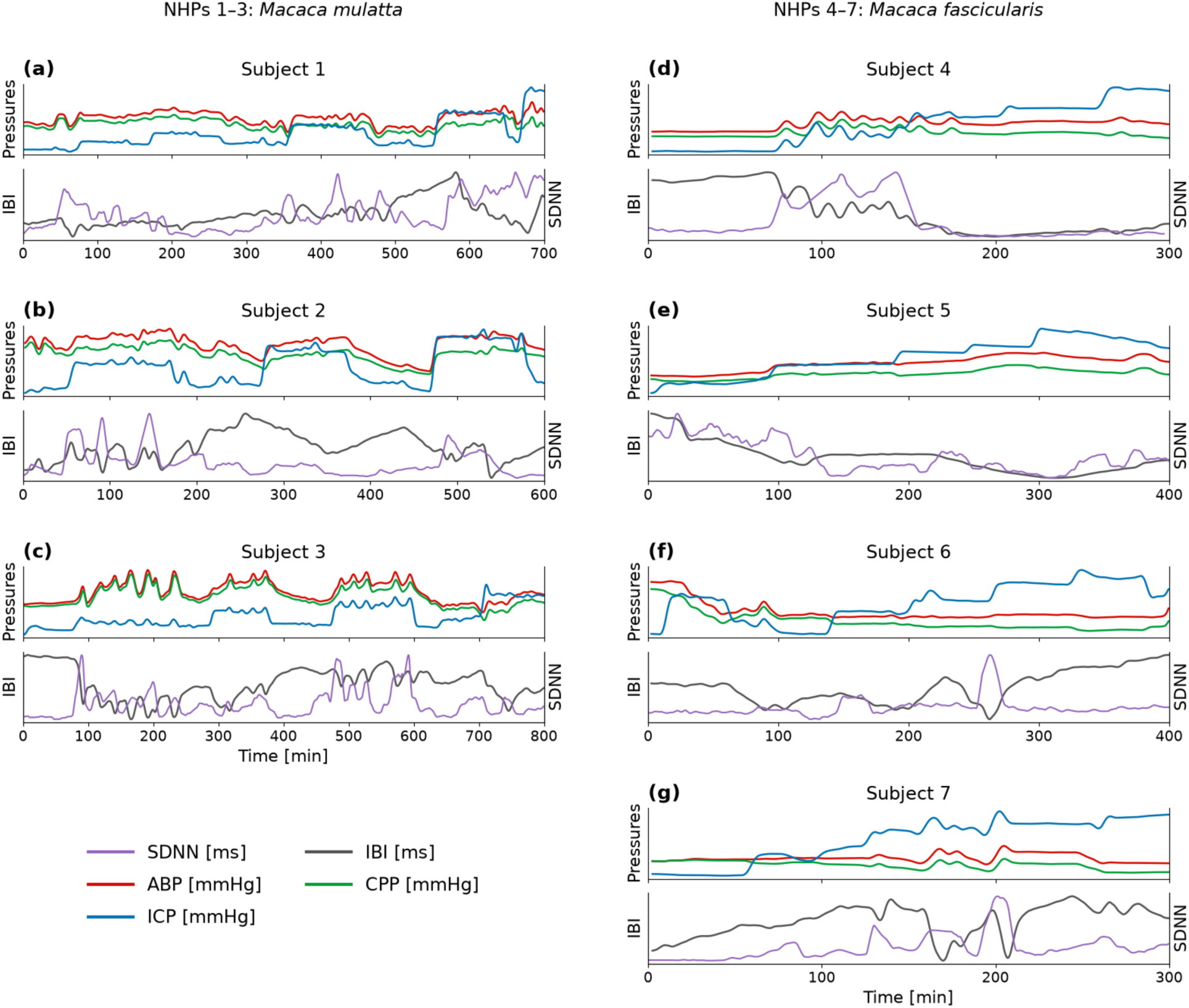
Physiological time series for each subject (n=7) with ABP, CPP and scaled ICP plotted on upper plot and IBI and SDNN in ms plotted on lower-level plot over the course of the recording.

To further examine how the relationship with HRV and ICP evolves over increasing ICP, Figure 3 plots SDNN, RMSSD, and PRx over ICP in increasing bins between 10-15mmHg increments and the corresponding CPP values for n=7 subjects pooled together. Individual subject averages are also plotted over the bars. Bin sizes were chosen to reflect the ICP increments used during the experiment where each increment aimed for a 10 mmHg increase as shown in the sample plot in Figure 1 while keeping the distribution of data as even as possible across both subjects and samples used in each bin, displayed in Figure 4.

**Figure 3.**
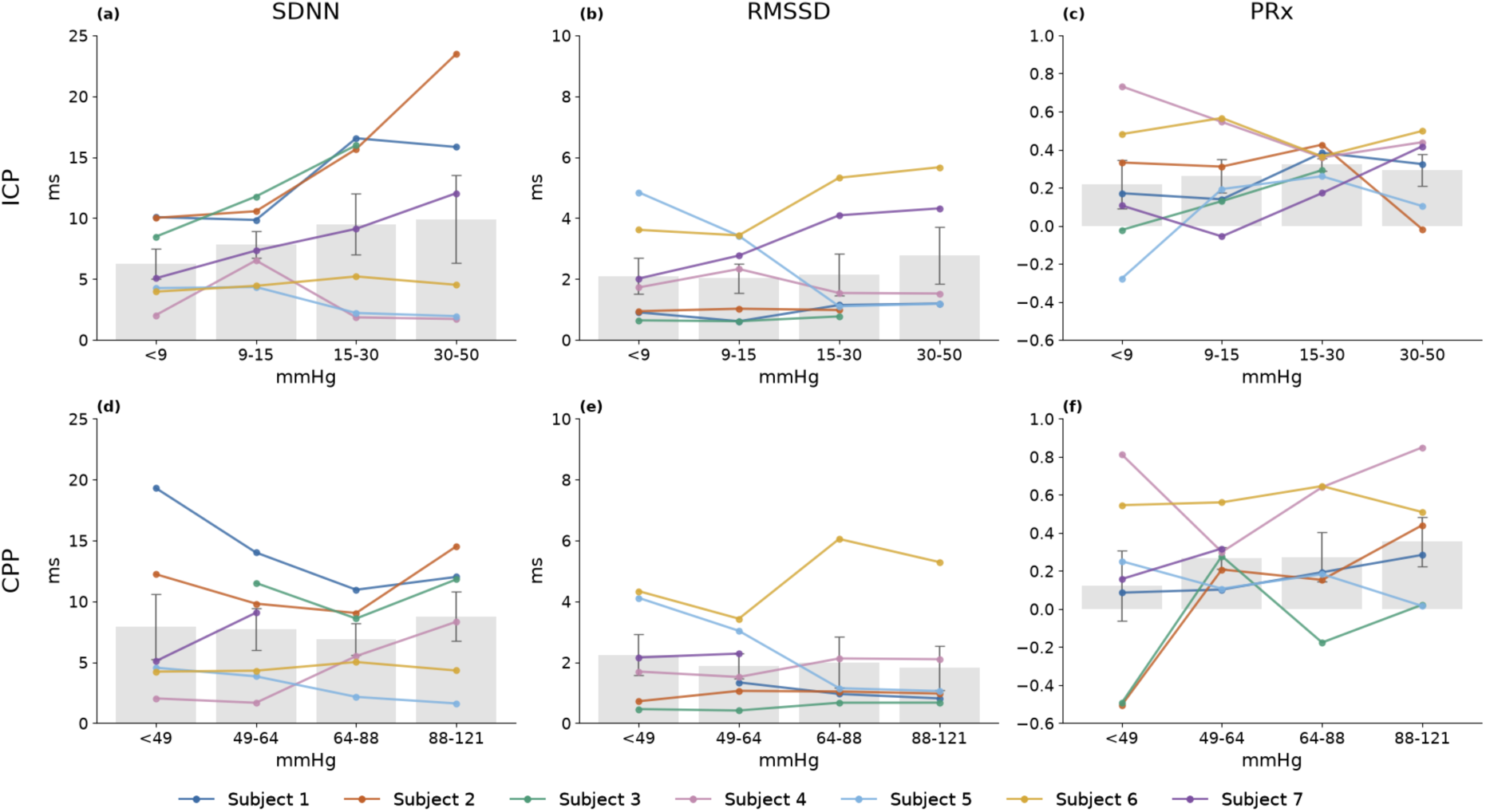
SDNN, RMSSD, and PRx across ICP bins (a-c) and corresponding CPP bins (d-f) pooled across n=7 subjects. Gray bars indicate pooled means for each bin with error bars indicating SEM. Colored lines show individual subject means. One-way repeated-measures ANOVA found no significant effect of ICP or CPP on SDNN, RMSSD, or PRx (p > 0.05).

**Figure 4.**
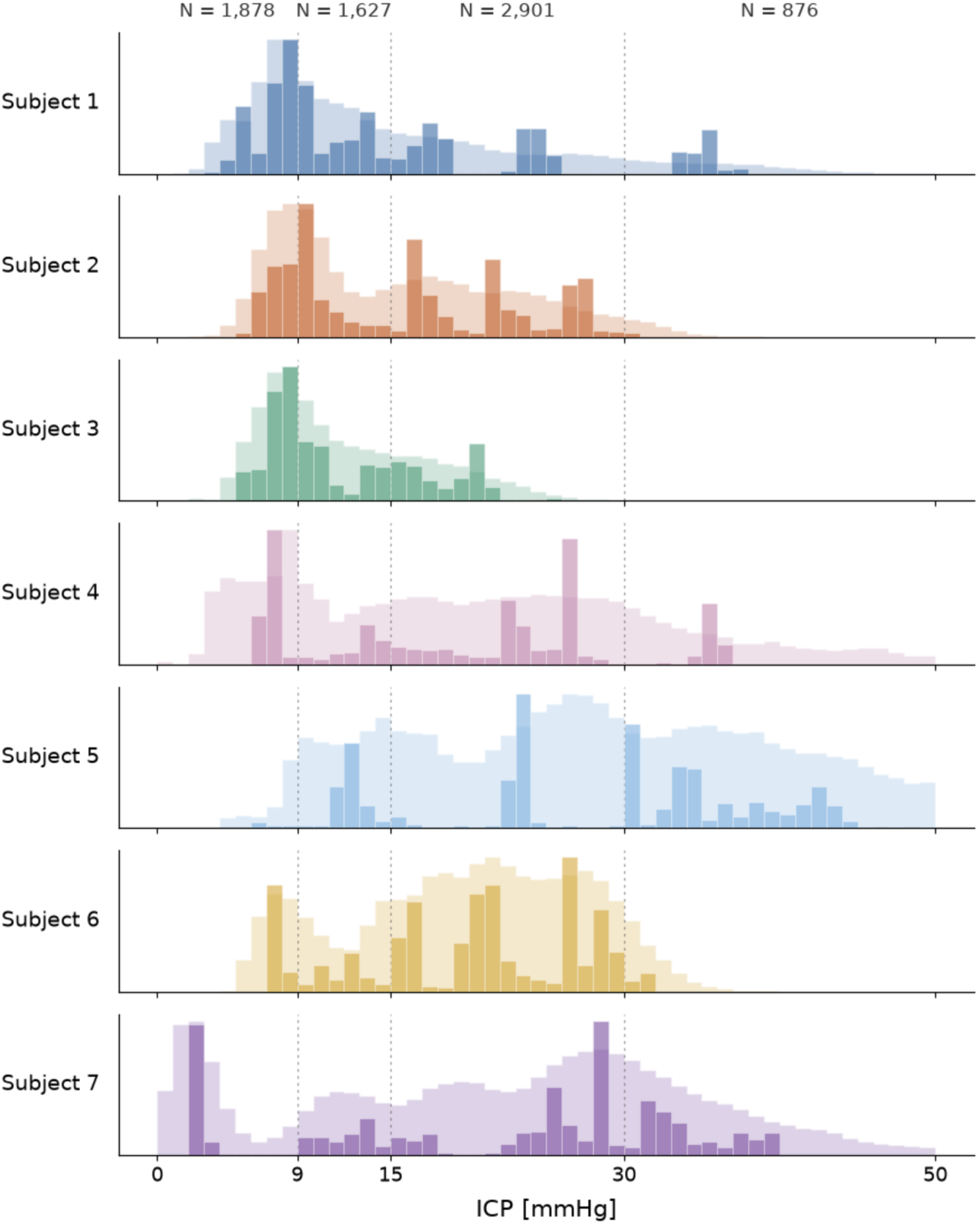
Distribution of ICP for each subject with relative frequency plotted on y-axes. Vertical dashed lines indicate bin boundaries used in Figures 3 and 6 with the number of samples N in each pooled bin indicated. Upper level of ICP values clipped at 50 mmHg to maintain even distribution between subjects. Darker bars indicate the ICP values averaged over the 60 second rolling window while the lighter bars show the full raw distribution before pre-processing.

In Figure 3a and 3b, RMSSD and SDNN increases with increasing ICP. The SDNN plotted over CPP demonstrates an inverse trend overall. The SDNN for all but subjects 4 and 5 increases over increasing ICP. Notably, subjects 2 and 3 had opposite CPP and ICP trends until the 15-mmHg bin at which point CPP and ICP had the same trend. The RMSSD increased overall for subjects 1, 4, 6, and 7. Subject 5’s RMSSD decreased while subjects 2 and 3 remained mostly flat. PRx, plotted in Figure 3c, does not reveal any distinct trends across subjects. However, the average is initially centered at .2 in the lowest ICP bin and increases up to the 15-20 mmHg bin size and then decreases again once ICP surpasses 30 mmHg. One-way repeated measures ANOVA was performed for each subplot with p=.05 with no significant effect of ICP on SDNN (ICP: F=1.774, p=0.195, n=6; CPP: F=0.713, p=0.563, n=5), RMSSD (ICP: F=0.040, p=0.989, n=5; CPP: F=0.169, p=0.915, n=5), or PRx (ICP: F=0.142, p=0.933, n=6; CPP: F=0.985, p=0.426, n=6).

Pairwise t-tests were performed with p=0.05 but also returned non-significant results. Because ICP levels varied across subjects, particularly at higher values, not every subject is represented in each ICP bin, which could have impacted between subjects’ results. T-statistics and p-values for all subjects are plotted in matrices in Figure 5. All subjects demonstrated a significant change in SDNN between the 9-15 to 15-30 mmHg ICP bin. However, the direction of the change was different, with subjects 4 and 5 showing a negative change. For RMSSD and PRx, 5 out of the 6 subjects demonstrated a significant change between the 9-15 to 15-30 mmHg ICP bin. These results highlight how although there are notable trends between ICP and HRV, these trends are highly sensitive to subject physiology and initial state and are not conclusive at the group level.

**Figure 5.**
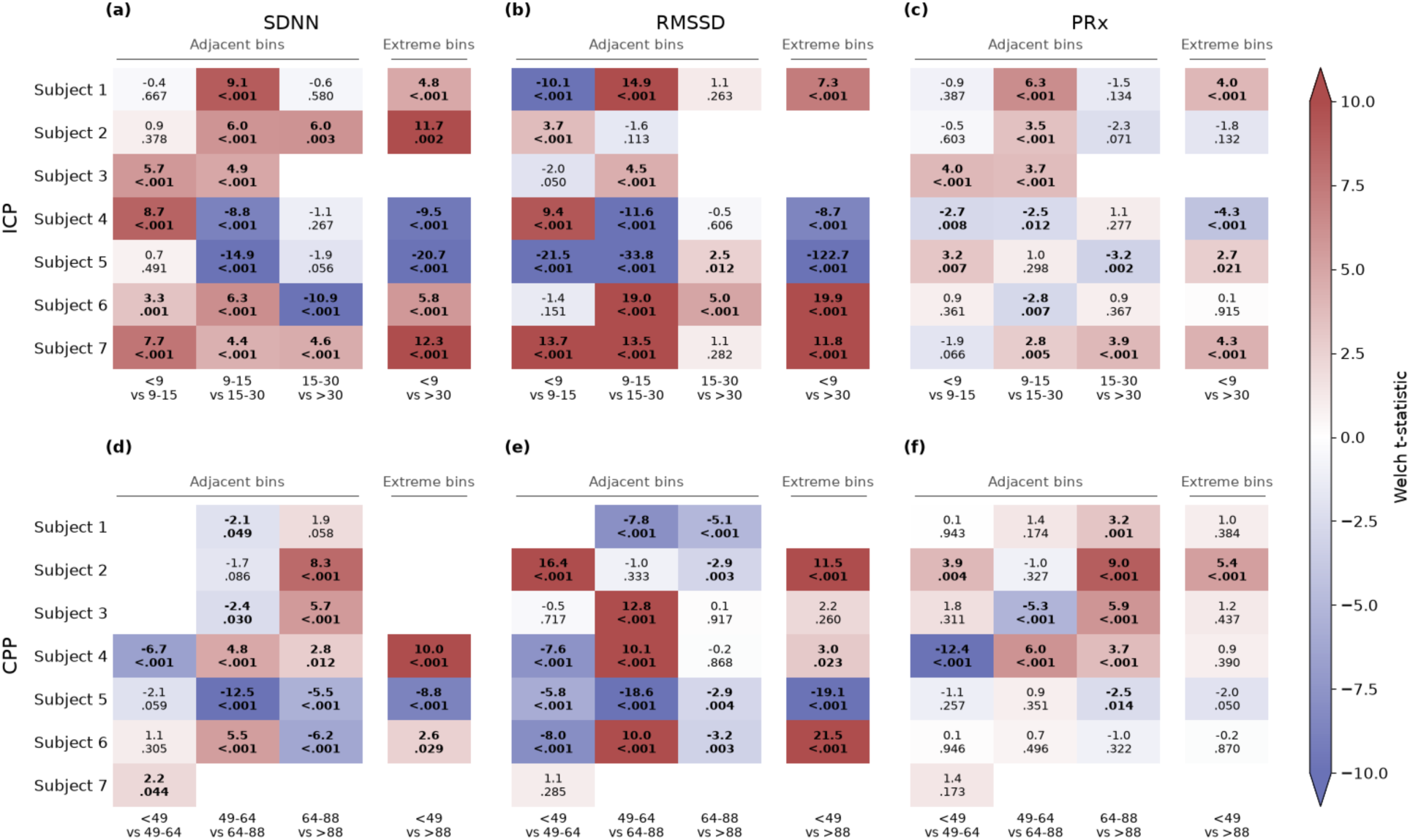
Subject-level pairwise comparisons of SDNN, RMSSD, and PRx between ICP bins (a-c) and CPP bins (d-f) for each subject. Cells show Welch’s t-statistic (top) and p-value (bottom) for each pair of bins. For each metric, the adjacent bins columns reflect tests between neighboring bins while the extreme bins column reflect a comparison between lowest and highest bins. Bold values indicate p <0.05 and white cells indicate pairs that could not be tested due to insufficient samples in bins.

### 3.2 Frequency Domain HRV Reveals No Clear Trend With ICP

Figure 6 plots frequency domain metrics of normalized LF, normalized HF, and LF/HF over the same ICP bins as described in Figure 3. Across the pooled subjects, LF/HF ratio decreases as ICP increases and increases as CPP increases. The normalized LF and HF values do not show a clear trend. One-way repeated measures ANOVA was performed for each subplot with p=.05. The ANOVA revealed no significant effect of ICP or CPP bins on the normalized LF (ICP: F=0.613, p=0.620, n=5; CPP: F=0.580, p=0.643, n=4), normalized HF (ICP: F=2.102, p=0.153, n=5; CPP: F=1.280, p=0.316, n=6), or LF/HF ratio (ICP: F=1.625, p=0.236, n=5; CPP: F=0.785, p=0.532, n=4). Pairwise t-tests found a significant difference between 49-64 mmHg and 64-88 mmHg CPP bin for the normalized LF (p=0.031, t=2.968). Despite no significant result at the group level, subjects 1, 2, and 4 show significant changes in the relationship between normalized LF and ICP between the 9-15 mmHg and 15-30 mmHg bins. The LF HRV for subjects 3 and 5 decreased as ICP increased while the normalized LF activity for subject 5 increased over increasing ICP before decreasing following the 15-30mmHg bin. The inverse relationship is demonstrated for the normalized HF for subjects 1, 2, and 5 with increasing HF activity before the transition between the 9-15mmHg and 15-30mmHg ICP bin. Subject 6 also demonstrates this relationship, though not with significance. In subjects 4 and 7, the normalized HF activity increased across increasing ICP before decreasing following the 15-30mmHg bin while subject 3 showed only increasing normalized HF over increasing ICP. This transition at the 15-30mmHg bin is also shown in the LF/HF ratio for subjects 1, 2, and 6 while subject 3 decreases, implying a larger role in the low frequency HRV activity. The LF/HF ratio value for subjects 4, 5, and 7 implies a dominance of the HF activity over the trial. For CPP, the inverse relationship is observed across subjects 1, 3, 4, and 5, with the change point occurring in CPP bins corresponding to the transition from the 9-15mmHg ICP bin and 15-30mmHg CPP bin. T-statistics and p-values for all subjects are plotted in matrices in Figure 7.

**Figure 6.**
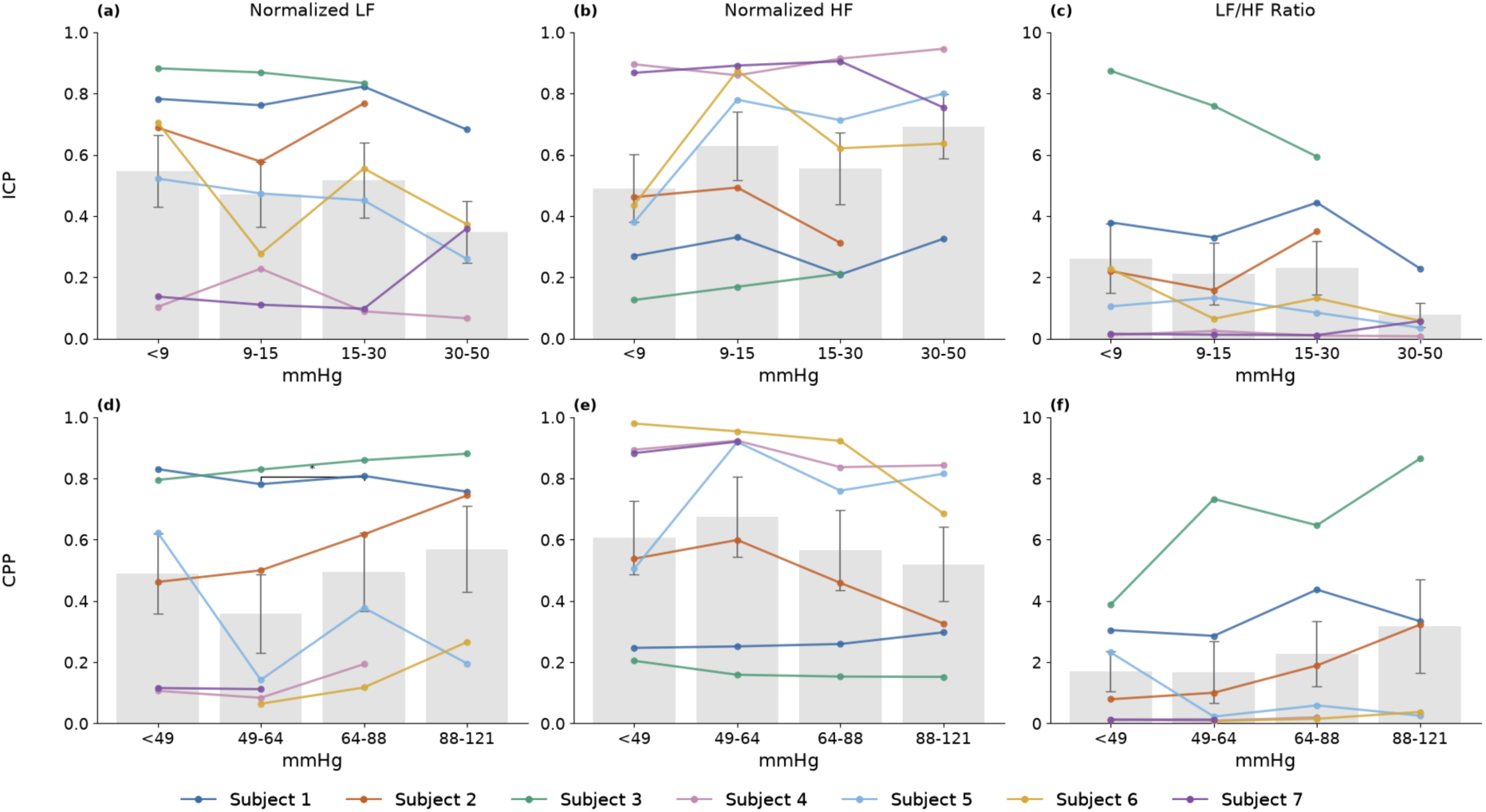
Normalized LF, Normalized HF, and LF/HF Ratio across ICP bins (a-c) and corresponding CPP bins (d-f) pooled across n=7 subjects. Gray bars indicate pooled means for each bin with error bars indicating SEM. Colored lines show individual subject means. One-way repeated-measures ANOVA found no significant effect of ICP or CPP on Normalized LF, Normalized HF, or LF/HF ratio (p > 0.05).

**Figure 7.**
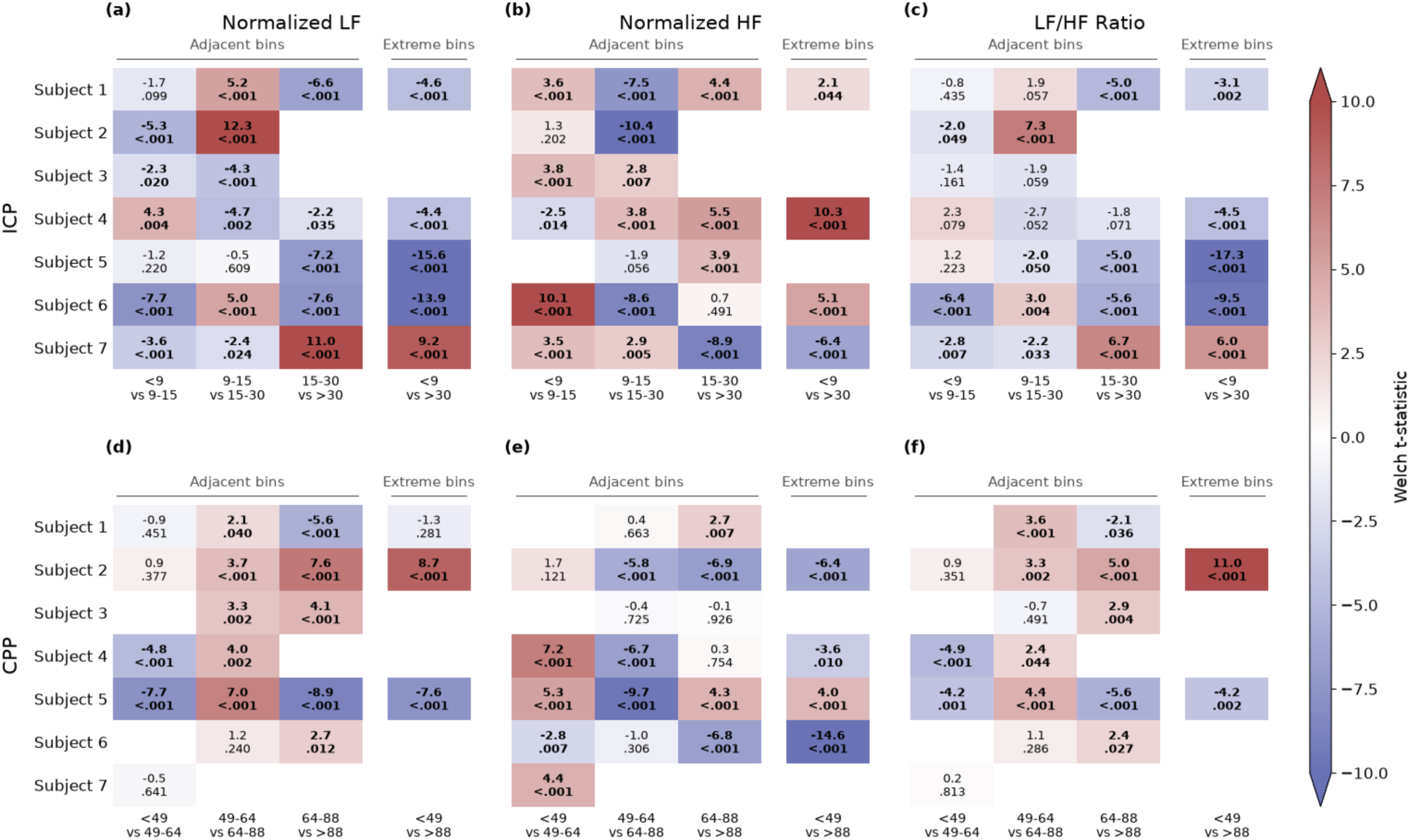
Subject-level pairwise comparisons of Normalized LF, Normalized HF, and LF/HF Ratio between ICP bins (a-c) and CPP bins (d-f) for each subject. Cells show Welch’s t-statistic (top) and p-value (bottom) for each pair of bins. For each metric, the adjacent bins columns reflect tests between neighboring bins while the extreme bins column reflect a comparison between lowest and highest bins. Bold values indicate p <0.05 and white cells indicate pairs that could not be tested due to insufficient samples in bins.

### 3.3 HRV Relates to Autoregulatory State

Figure 3c and 3f shows that across pooled subjects, PRx passes the clinical threshold at the same ICP bins at which the relationship between HRV and ICP begins to change, suggesting an influence of CAR on HRV over increasing ICP. In healthy conditions, a drop in blood pressure will trigger dilation of blood vessels to maintain stable blood flow and an increase in blood pressure will trigger constriction of blood vessels, reflecting an inverse relationship between ICP and ABP. In dysfunctional conditions, however, ICP moves with blood pressure changes, indicating that the blood vessels can no longer react to changes in pressure and blood flow. Because raised ICP can obstruct cerebral blood flow and thus impact CAR an association between CAR and HRV may indirectly reflect ICP increases. The influence of CAR on HRV is investigated in Figure 8, which plots time- and frequency-domain HRV metrics before and after the PRx threshold. A paired t-test with significance p<0.05 was performed on the group average of the subjects, reaching statistical significance for SDNN (p= 0.024, t= 3.00), but not for RMSSD (p= 0.712, t= 0.388), Normalized LF (p= 0.130, t= 1.757), Normalized HF (p=0.129, t= -1.757), or the LF/HF Ratio (p=0.425, t=.856). On the subject level, significant differences are observed for most subjects for SDNN and frequency metrics. P values from Welch’s t-test with significance p<0.05 for subject differences are reported in Table 1. Notably, for SDNN and Normalized LF, all subjects except for subject 5 increased between the two PRx groups. For Normalized HF, all subjects except for subject 5 and subject 3 decreased between the two PRx groups.

**Figure 8.**
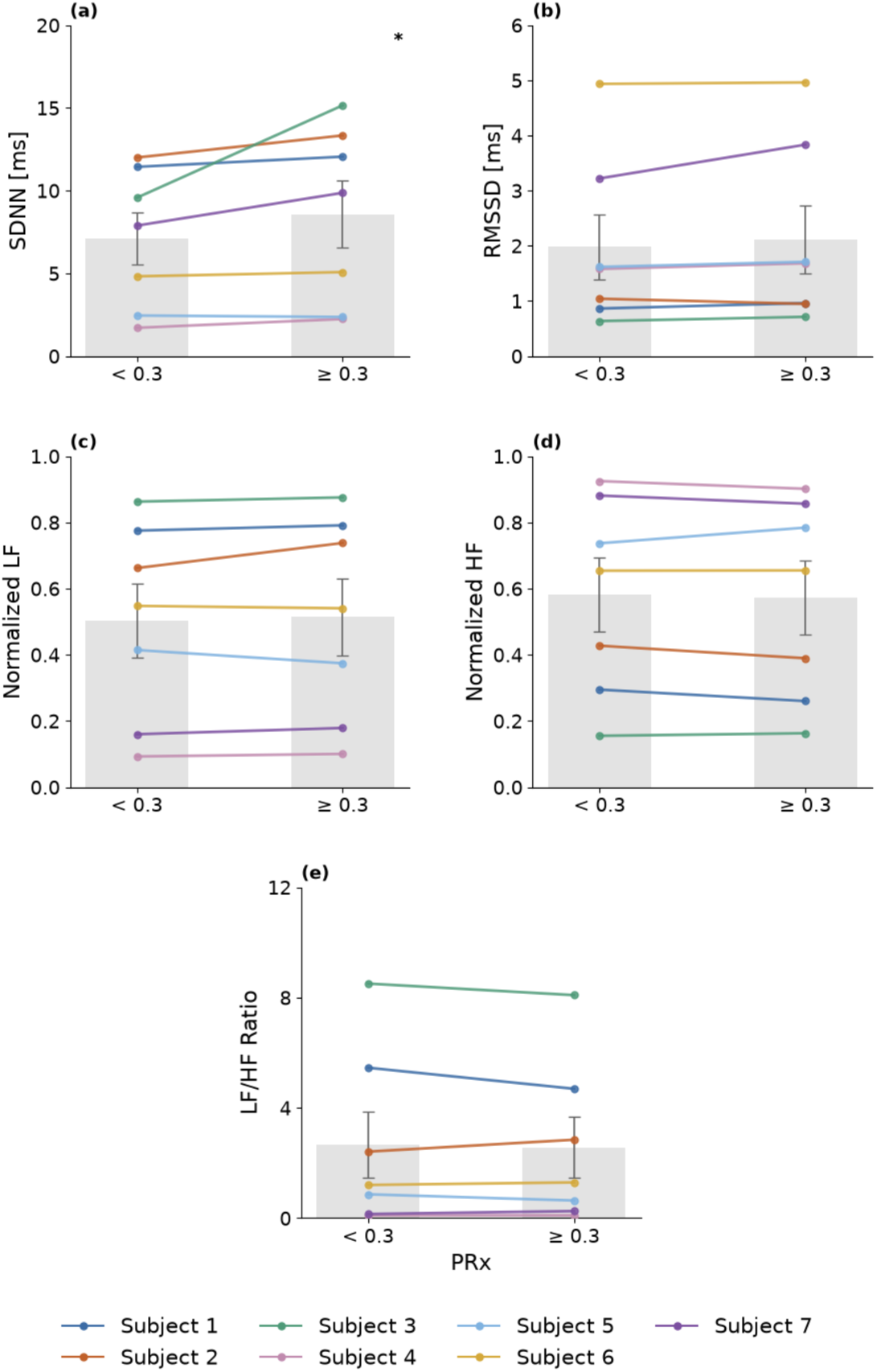
Time- and frequency-domain HRV metrics by PRx group (< 0.3 vs ≥ 0.3) pooled across n=7 subjects. Gray bars indicate pooled means for each bin with error bars indicating SEM. Colored lines show individual subject means. Asterisks denote a significant difference between groups (paired t-test on subject means, * p < 0.05). (a) SDNN, (b) RMSSD, (c) Normalized LF, (d) Normalized HF, and (e) LF/HF ratio.

**Table 1.** Welch’s t-test results per subject comparing HRV metric before and after critical PRx threshold.

| **Subject** | **SDNN** |  | **RMSSD** |  | **Normalized LF** |  | **Normalized HF** |  | **LF/HF Ratio** |  |
|---|---|---|---|---|---|---|---|---|---|---|
|  | t | p | t | p | t | p | t | p | t | p |
| 1 | 4.610 | <0.001 | 4.580 | <0.001 | 3.752 | <0.001 | -3.752 | <0.001 | 2.242 | 0.026 |
| 2 | 3.405 | 0.001 | -2.461 | 0.014 | 6.943 | <0.001 | -6.943 | <.0001 | 2.925 | 0.003 |
| 3 | 7.408 | <.001 | 2.493 | 0.013 | -1.567 | 0.118 | 1.567 | 0.118 | -2.454 | 0.014 |
| 4 | 2.810 | 0.006 | 1.087 | 0.283 | 3.174 | 0.003 | -3.174 | 0.003 | 3.433 | 0.001 |
| 5 | -0.694 | 0.488 | 3.198 | 0.002 | -3.402 | 0.001 | 3.402 | <0.001 | -1.092 | 0.276 |
| 6 | 2.816 | 0.005 | -5.304 | <0.001 | 3.543 | <0.001 | -3.543 | <0.001 | 5.455 | <0.001 |
| 7 | 3.002 | <0.001 | 3.739 | <0.001 | 4.024 | <0.001 | -4.024 | <0.001 | 1.604 | 0.110 |

## 4. Discussion

Our results directly highlight that HRV can change for a variety of reasons, and these changes are not always directly correlated with ICP. Although HRV trends followed ICP increases, these trends were not significant at the group level. Figures 3-7 demonstrate significant subject level differences between ICP bins in both time and frequency metrics, but no clear linear relationship between changes in ICP and HRV at the group level. Although no significant group level trends were observed, individual subjects’ HRV had distinct progressions up to a threshold of ICP that then reversed direction for the remainder of the bins. Interestingly, in both the time and frequency domain metrics, this change in trend appeared between the 9-15mmHg and 15-30mmHg bins across subjects, just below the ICP threshold where mortality and unfavorable outcomes are typically observed (Rapp *et al* 2025). While the precise threshold is controversial, the existence of a patient dependent ICP threshold is well reported (Riparbelli *et al* 2024, Lazaridis *et al* 2020). These results are also consistent with previous reports that necessitate interpreting HRV changes relative to a patient’s baseline rather than pre-determined linear relationships (Ghazzal *et al* 2026). This reversal in relationship is also observed when comparing between ICP and CPP trends. At CPP levels corresponding to low ICP bins, the HRV vs ICP and HRV vs CPP had opposite trends as seen in Figures 3a, b, d, and e and in Figure 6. As ICP increased, the HRV vs ICP and HRV vs CPP began to follow the same trend. This can be observed for subjects in all HRV metrics.

The observed change in the relationship between CPP and ICP is consistent with the literature that describes an inverse relationship when there is regular autonomic functioning and a disruption of this relationship in physiologically abnormal cases (Czosnyka *et al* 1997). Such an ICP threshold is clearly demonstrated in Figure 2, which shows how a spike in ICP triggers a change in the relationship between the IBI, CPP, and ABP. Subjects show that before a critical ICP value, IBI does not track ABP and CPP changes, indicating normal autoregulatory functioning. Following the spike, the IBI begins to oscillate with the pressure variables. At ICP elevations ABP, ICP, and IBI all oscillate together, but fall out of sync as ICP stabilizes. This directly illustrates in time how the heart tracks elevations in ICP that impact autoregulation. However, these observed changes could not be generalized to subjects or particular ICP levels, which reflects the growing skepticism about the existence of discrete ICP threshold (Riparbelli *et al* 2024).

In fact, the only significant group level relationship observed was the correlation between SDNN and PRx in Figure 8a. This demonstrates that when using HRV as a diagnostic measure in TBIs where ICP is elevated, it is far more reflective of changes in CAR rather than changes in ICP. An increasing PRx in response to increased ICP in the case of TBI has been observed in multiple clinical studies, along with reports of increasing amplitude of the cardiac pulse in ICP. Donnelly et al. note that in some patients, once ICP reached a critically high threshold, the relationship between the cardiac pulse and ICP switched, which can be indicative of cerebrovascular collapse (Donnelly *et al* 2019, 2017) which can be explained by the notion that an increase in ICP past a critical point can cause venous compression such that ICP and CPP have a positive relationship, causing decrease in perfusion (Pinto and Adeyinka 2025). Observations from Figures 2, 3 and 5, closely align with Donnelly’s observations, showing a coupling between heart dynamics and ICP as ICP increases as well as similar switching behavior past a critical threshold. Figure 8, which plots HRV metrics as a function of PRx before and after a threshold of 0.3, shows directly how HRV metrics relate to the state of autoregulation with a significant change in the HRV after the critical PRx threshold. ICP and cardiac pulses may be more coupled as ICP increases due to increased attempts of compensatory reflexes, such as the Cushing reflex, to restore stable perfusion (Dinallo and Waseem 2023). It is possible that at lower ICP levels, there need not be tight coupling of ICP and HRV because the brain can maintain perfusion through the normal vascular response. However, as ICP reaches dangerous levels, physiological reflexes will have to engage more to maintain stable perfusion. This can lead to further increase in ICP as an acute process.

This study’s findings also speak to the controversies concerning frequency domain metrics and autonomic state. Frequency domain HRV metrics may reflect the changing state of cerebral autoregulation through ICP fluctuations. Lower LF/HF ratio has been shown to correspond to worse patient outcomes and cerebral functioning with an increased ICP potentially leading to cerebral ischemia of neurons that are responsible for mediating circulation in the brain and cardiovascular reflexes. It has also been postulated that ICP and sympathetic activity are closely connected with a study by Schmidt et al. showing that ICP increases trigger sympathetic activity and a vagal withdrawal as ICP rises (Schmidt *et al* 2018, Biswas *et al* 2000). The HF HRV results reflect this, with a drop in normalized HF power after critically elevated ICP in Figure 6b and a drop in the normalized HF power between PRx threshold in Figure 8. The switching behaviour observed in Figure 6b could be attributed to collapsing cerebral vasculature as ICP approaches lethal levels. However, no group level significance emerged from this analysis. Further, many groups report conflicting evidence on how frequency domain HRV and autonomic signals are related, leaving the extent to which the frequency domain HRV reflects the autonomic nervous system activity a topic of debate (Billman 2013, Reyes del Paso *et al* 2013, Hayano and Yuda 2019). While Schmidt et al. claim that changes in LF reflect sympathetic activity, other studies claim that that sympathetic activation cannot be disentangled from the IBI power spectrum and most directly reflects baroreflex function (Goldstein *et al* 2011). A lower LF power may reflect diminished slow oscillations, indicating a weakened baroreflex response. Results in Figure 6 show drops in the normalized LF and LF/HF power as ICP increases, perhaps reflecting failure of the baroreflex as ICP increases. Figure 8, however, shows an increase in normalized LF and the LF/HF ratio between the PRx bins. The inconsistencies in these findings underscore a need to be wary when interpreting frequency components of HRV (Reyes del Paso *et al* 2013). Further, while these thresholds can be useful for patient monitoring, they may not reflect the full story, with many studies pointing out that assumptions of cerebral hemodynamic do not hold in cases of autoregulatory breakdown. This highlights the importance of interpreting these variables carefully and relative to subject physiology (Stroh *et al* 2024, Lazaridis *et al* 2020).

## 5. Limitations

Key limitations of the study include the small subject pool and inter-species differences. Differences in measurement techniques between species may have also impacted results as HRV is highly sensitive to small deviations in the beat interval. Inconsistencies between the ICP spread between subjects may have affected results as well. Future studies with a larger and more homogeneous subject pool and a more even spread of ICP values should be conducted. Further, the relationship between autoregulation and cerebral vasculature could be further probed by inducing causal changes to other pressures in the vasculature.

## 6. Conclusions

The purpose of this study was to characterize HRV changes with changing ICP to better understand autonomic function and autoregulation in cases of high ICP. The present study found that there is a positive relationship between time domain measurements of HRV and PRx and confirmed that autoregulation degrades with increasing ICP. Results show no significant relationship between HRV metrics and ICP, but that increases in ICP can lead to dysfunctions in CAR and changes in autonomic control. This suggests that HRV could be a valuable biomarker for an ICP threshold at which autoregulation is not intact. While the precise physiological underpinnings of these changes remain elusive, results still indicate a potential clinical biomarker for autoregulatory changes and an indirect measure of elevated ICP in TBI that could be determined on a subject level basis. Further, the lack of discernible trends in this work reflects broader discrepancies in the literature. Reporting such findings is crucial for eventually teasing apart the mechanisms underlying these complex physiological interactions.

## Acknowledgements

The authors thank Samantha Schmitt, Matthew A. Smith, and Deepa Issar for their contributions to data collection and valuable scientific discussions.

## Notes

### Competing Interest Statement

The authors have declared no competing interest.

### Funding

- National Institutes of Health: R21-EB024675
- American Heart Association: 17SDG33700047
- Pennsylvania Institute of Technology
- Center for Machine Learning and Health, Carnegie Mellon University

